## Supplemental Figures for "Canonical WNT signalling governs *Echinococcus* metacestode development"

### S1 Figure

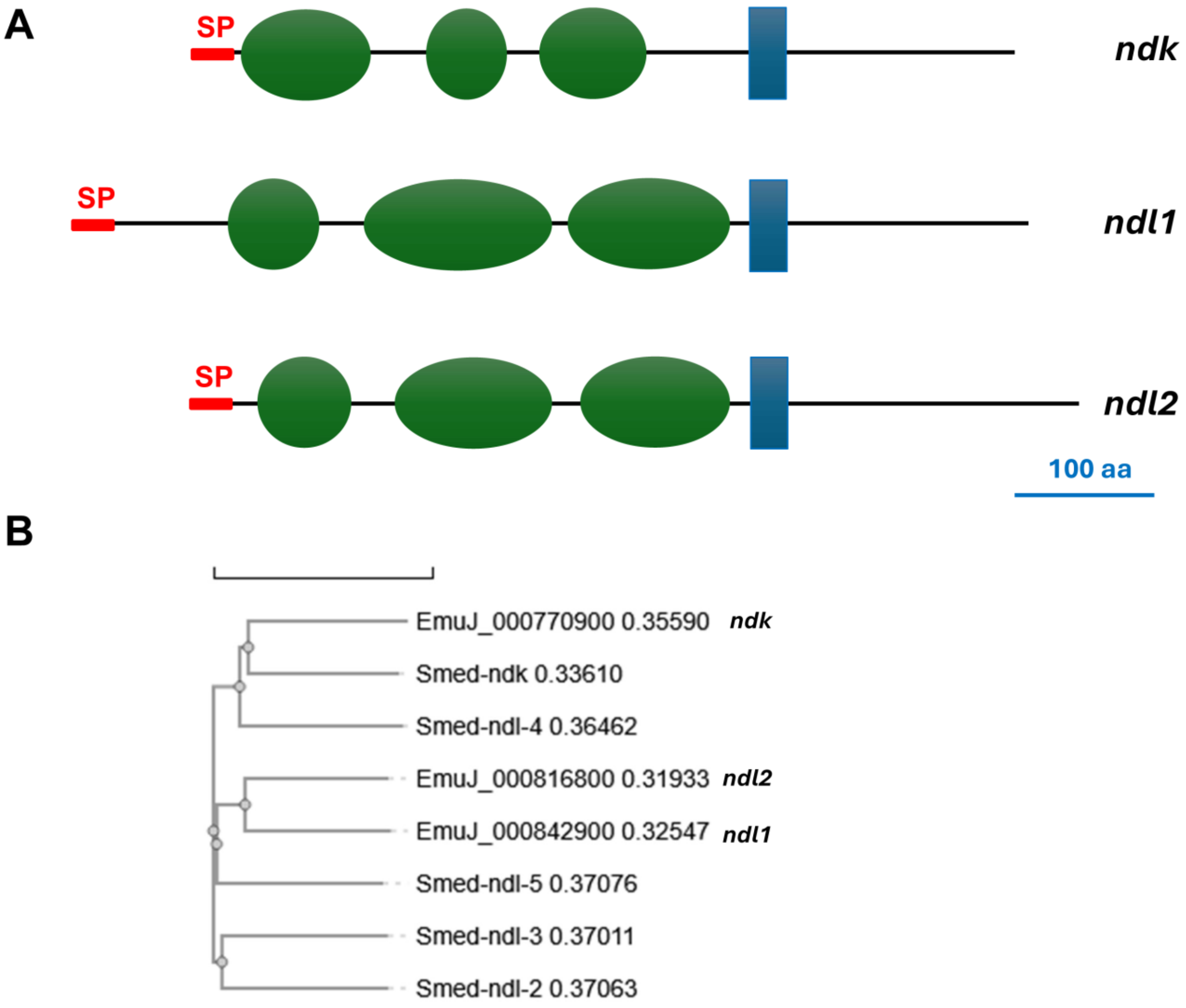


**S1 Figure. Structure and homologies of *Echinococcus ndk* and *ndl* (*ndk*-like) factors.** (A) Domain structure of *E. multilocularis ndk*, *ndl1*, and *ndl2* (as indicated). Shown are transmembrane domains (blue), Ig-Domains (green), and signal peptides (red). Please note that all three proteins display the characteristic 3 extracellular Ig-domains, but no intracellular kinase domain (as in FGF receptors). Size bar indicates 100 amino acids. (B) Phylogenetic analysis of *Echinococcus ndk*, *ndl1*, and *ndl2* with *Schmidtea mediterranea ndk* and *ndl's*. Depicted are the *Echinococcus* factors with gene ID and name as well as all *Schmidtea ndk*-like factors.

S2 Figure

A

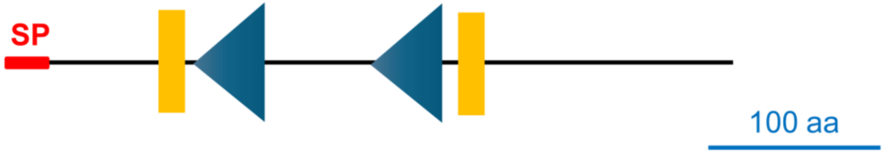


B

```
FST_HUMAN       MVRARHQPGGLCLLLLLLCQFMEDRSAQAGNCWLRQAKNGRCQVLY-KTELSKEECCSTG      59
FST_ECHMU       ---------MVPLLIFVLQALLVFTVSDAGSCFRIRLPNGRCHELLQRNYITRSECCRLG      51
FST_SCHMA       ----------------------------------------------MEESISQLECCKRG      14
                                                               .  ::: ***  *

FST_HUMAN       RLSTSWTEEDVNDNTLFKWMIFNGGAPNCIPC------KETCENVDCGPGKKCRMNKKNK      113
FST_ECHMU       GFYVSGHL--TEPQFVRDLLLRRTGIANCHYSCQRGFHNESCARVNCPDGFRCEVQN-HE      108
FST_SCHMA       GFYLNRQI--TQQEYLTDHLLFESGTPNCVQACNEELP-STCKGYVCPEGHYCRMIN-NE      70
                 :  .     .: : : . :: . *  **  .       .:*    *  *  *.: : ::

FST_HUMAN       PRCVCAPDCSNITW-KGPVCGLDGKTYRNECALLKARCKEQPELEVQYQGR-----C---      164
FST_ECHMU       AVCACSSICIVEDFESGPVCTTDFRQFRNRCSFLKERCRNLDTLLVEMECPPADHVCPYR      168
FST_SCHMA       PKCKCRLQCNTADYLTGPLCTTNLRRFHNQCHLKQARCKSPKESLEVIPCPDEGLRCSDL      130
                  * *   *    : .**:*  : : ::*.* : : **:.                *

FST_HUMAN       ------------------------------------------------------------      164
FST_ECHMU       QNTS--------------------------------------------------------      172
FST_SCHMA       SILSEKNLNLNSLFKTLYAEYEERKGVPSLPSSSSLPSSTSTTKSSLSSTDNQEYTDNED      190

FST_HUMAN       -----------------------------------KKTCRDVFCPGSSTCVVDQTNNA-Y      188
FST_ECHMU       ------------------------HLE------DHDYAYPVRICPPNKVCIVRQYSGRAS      202
FST_SCHMA       DNVIIMNRRTSTDLKTYIDKMNDESIFDEKSPYSNDELSKELICNANEYCLLRQFDGRPS      250
                                                   .      :*  .. *:: * ..

FST_HUMAN       CVT-------------------CNRICPEPASSEQYLCGNDGVTYSSACHLRKATCLLGR      229
FST_ECHMU       CEDPNWLGRKHIESAFDRVAVSNLDSCANS-PYGSPICGTDNNTYNNQCHLRKASIQKGL      261
FST_SCHMA       CEYWN------------EQSWQTLNNCPES-TFNEPICSTDNQTFYNSCDLKLTGLKNQL      297
                *                         * :     . :*..*. *: . *.*: :

FST_HUMAN       SIGLAYEGKCIKAKSCEDIQCTGGK-KCLWDFKVGRGRCSLCDELCPDSKSDEPVCASDN      288
FST_ECHMU       EIRIAYLGPCRKNATCDQIRCQSPHMTCVPHVRTGQPVCLDCTDLPPNCNPRVFVRINPA      321
FST_SCHMA       EVRIAYKGECRSKVTCSDIQCPRPDMRCSPHHLTGQPICMDCAKLPLDCNASFRNGQETL      357
                .: :** * * .  :*.:*:*   .  *  .  .*:  *  * .*  :.:       .

FST_HUMAN       ATYASECAMKEAACSSGVLLEVKHSGSCNSISEDTEEEEEDEDQDYSFPISSILEW----      344
FST_ECHMU       EI---------AEMTLFEMFKEAKQ-----NF---GDPRWHGDVFTXWLASGMWKWKKVI      364
FST_SCHMA       TSSTSSSSSSSSSTGVLGKRKTQ-----IF---GDPVWKTPVQTY-------GWPVVC      402
                           :  :   : :   .           :   .              *

FST_HUMAN       ------------------------------------------------------------      344
FST_ECHMU       PQCLLPTRLQLYRQTLRRPCLNWILSSRVISELSIVR----------PQEQLKRYFILKK      414
FST_SCHMA       GS---------NGKTYRNTCFLHVVNCLSDKFVGLKKPEFCSETIPSKHDDWRRHFGK--      451

FST_HUMAN       ----------------------------------------    344
FST_ECHMU       ERTSEVKSITTIITTTFVTLTLASPHCCLSCALMHATQDP    454
FST_SCHMA       --------YSNL----F-----------------------    456
```

**S2 Figure. Structure and homologies of *Echinococcus fst*.** (A) Domain structure of *E. multilocularis* FST. Shown are characteristic follistatin N-terminal domains (FOLN, yellow), Kazal-type serine protease inhibitor domains (blue), and signal peptide (red). Size bar indicates 100 amino acids. (B) Amino acid sequence comparison betwen *E. multilocularis* FST (ECHMU), human FST (HUMAN), and *Schmidtea mediterranea* FST (SCHMA). Sites of perfect alignment (*) as well as groups of strong (:) or weak (.) similarity are marked below the sequences.

### S3 Figure

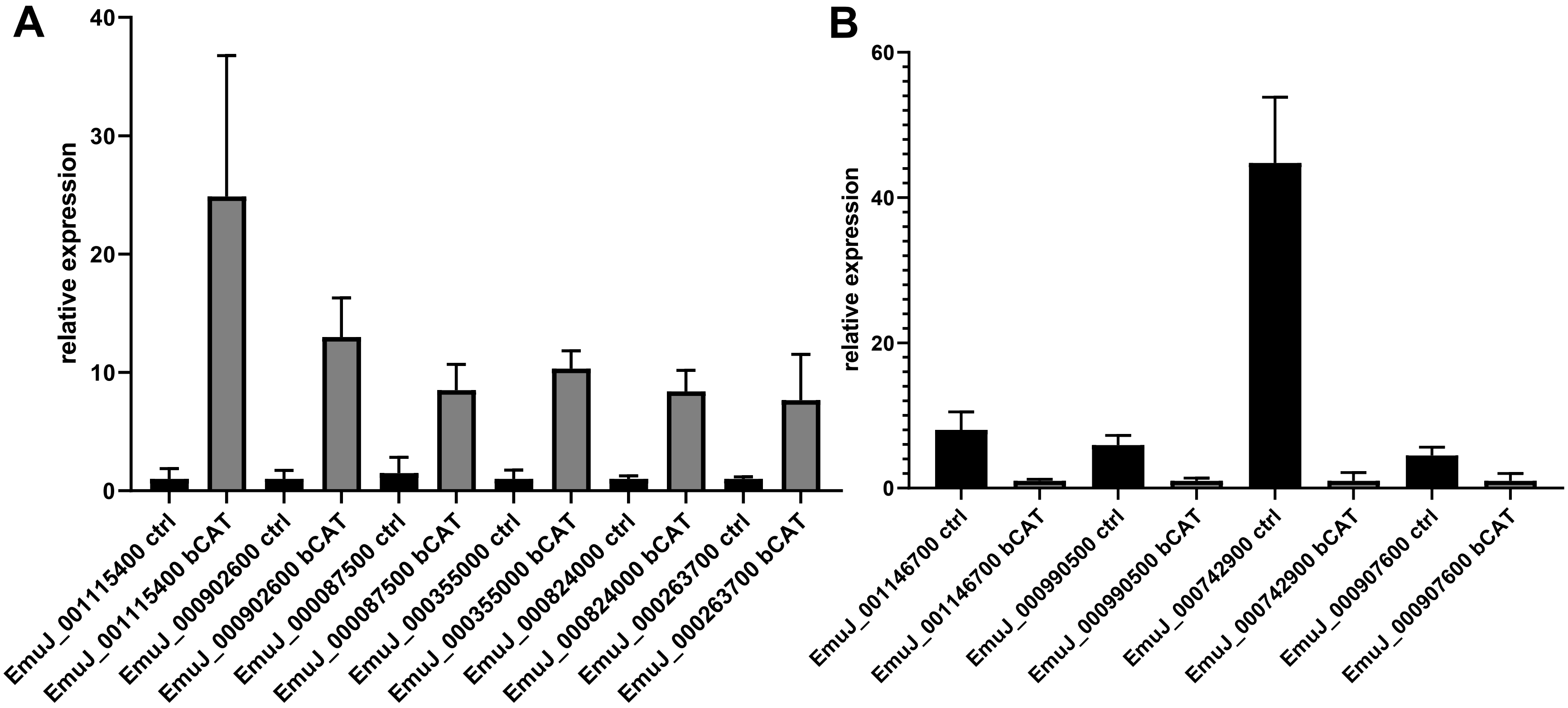


**S3 Figure. qRT- PCR analysis of gene expression after siPOOL RNAi.** A selection of genes that were significantly upregulated (A) or downregulated (B) after RNAi with siRNAs were analysed by qRT-PCR in cell cultures after siPOOL RNAi. Shown is relative expression in comparison to control gene *elp* (EmuJ_000485800). Indicated are gene IDs in control cultures (ctrl) and RNAi cultures (bCAT). Error bars indicate SD of three technical replicates.

### S4 Figure

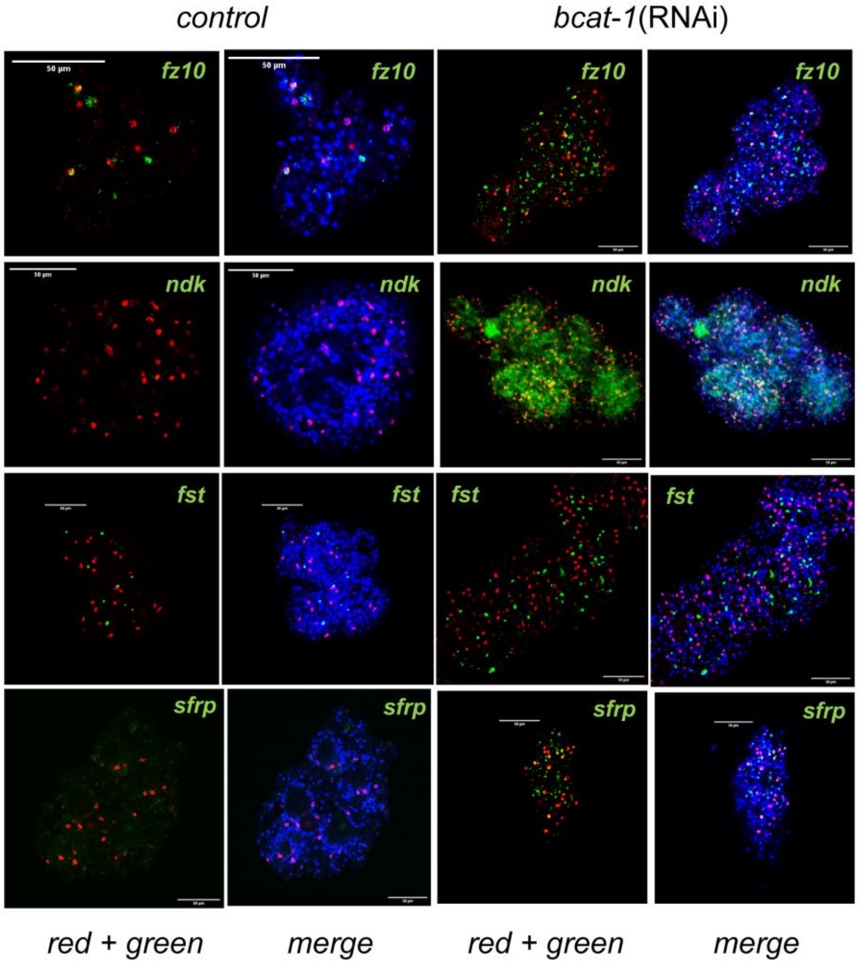


**S4 Figure. Induction of anterior markers in *bcat-1*(RNAi) culture aggregates.** WISH has been carried out on control RNAi cultures and bcat-1(RNAi) cultures (as indicated) for fz10, ndk, fst, and sfrp (from top to bottom). Shown are representative pictures for single confocal slices for red and green channel (red, Edu, proliferative stem cells; green, gene specific probe) and merge pictures also including third channel (blue, DAPI, nuclei) as indicaterd. Size bar represents 50 μm in all slides.

### S5 Figure

## A

*Echinococcus multilocularis* MUC2

```
MYFTQANDSASSDASAT                                                         17

PTTRSTPKVTNPNLTTTGSTPTNPTTPKVTKPTVSTTASPPTSRTTPKVTKPTLTTTRAT              77

PTTRSTPKVTNPNLTTTGSSPTTRTTPQLTKPTATTTKATPTTRTTPKVTKPTLATTGST             137

PTTRTTPQLTKPTATTTSSTPTTQTTQITPKVTKPAATTTKATPTTRTTPKVTKPTDSTT             197

SSTPSNRTTPKLTKPTVSTTGSTTSTSTTPDISTPAATTTTATCPTCPSSSSSSPNTTAT             257

ETSAPIPPSICFWCLSPVWIAVIVVCVVLVCAVLVVICIVKFCCCRSDRAGRPGRGGQNS             317

NLLVNVNQV                                                                326
```

## B

```
EmMUC2  18   PTTRSTPKVTNPNLTTTGSTPTNPTTPKVTKPTVSTTASPPTSRTTPKVTKPTLTTTRAT  77
             P TR  P  T P      S P     P  T P      S P +R  P  T P      +
HsMUC1  222  PDTRPAPGSTAPPAHGVTSAPDTRPAPGSTAPPAHGVTSAPDTRPAPGSTAPPAHGVTSA  281

EmMUC2  78   PTTRSTPKVTNPNLTTTGSSPTTRTTPQLTKPTATTTKATPTTRTTPKVTKPTLATTGST  137
             P TR  P  T P      S+P TR  P  T P A    + P TR  P  T P      S
HsMUC1  282  PDTRPAPGSTAPPAHGVTSAPDTRPAPGSTAPPAHGVTSAPDTRPAPGSTAPPAHGVTSA  341

EmMUC2  138  PTTRTTPQLTKPTATTTSSTPTTQTTQITPKVTKPAATTTKATPTTRTTPKVTKPTDSTT  197
             P TR  P  T P A   +S P T+     P  T P A    + P TR  P  T P
HsMUC1  342  PDTRPAPGSTAPPAHGVTSAPDTRP---APGSTAPPAHGVTSAPDTRPAPGSTAPPAHGV  398

EmMUC2  198  SSTPSNRTTPKLTKP  212
             +S P  R  P  T P
HsMUC1  399  TSAPDTRPAPGSTAP  413
```

**S5 Figure. Structural features of *Echinococcus* MUC2.** (A) Amino acid (aa) sequence of *E. multilocularis* MUC2 showing several highly similar, threonine-rich repeats between aa 18 and 257. Numbers to the right indicate MUC2 amino acids. A predicted transmembrane region is underlined. (B) Amino acid sequence comparison betwen repeat domains of MUC2 and human mucin MUC1 to which it shows highest homologies in SwissProt. Identical residues between both sequences are shown in interline. Biochemically similar residues are indicated by ‚+'.

### S6 Figure

**A**

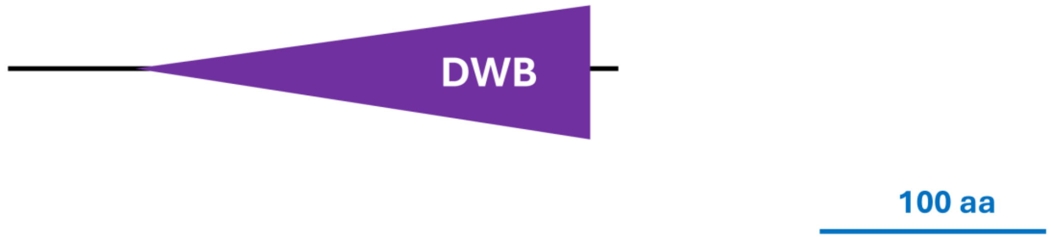


**B**

```
EmSmadF         KPWSHVSYWECNRHIGHKWLPVITSALVVLNEEHYSVSAATAENTSTHAVWSQQGRLCLD 60
SMAD6_HUMAN     SHWCSVAYWEHRTRVGRLYAVYDQAVSI-------------------FYDLPQGSGFCLG 41
SMAD7_HUMAN     SHWCVVAYWEEKTRVGRLYCVQEPSLDI-------------------FYDLPQGNGFCLG 41
                . *. *:*** . ::*: :     :  :                   .    * . :**.

EmSmadF         RLVRHFTSPRAFSEESHPSRKIYHPSILHSSSPTKQRRRLSHEGISLLLLPKGQILLTNK 120
SMAD6_HUMAN     QLNLEQRSE-------------------------SVRRTRSKIGFGILLSKEPDGVWAYN 76
SMAD7_HUMAN     QLNSDNKSQ-------------------------LVQKVRSKIGCGIQLTREVDGVWVYN 76
                :*   .  *                           ::  *: * .: *  : : : . :

EmSmadF         TLTTPIFVASPCFVQPGDLIAGDWPVYRVAPACSLVVFDTRIYED---RLTEAGKYTPWP 177
SMAD6_HUMAN     RGEHPIFVNSPTLDAPGGRA---LVVRKVPPGYSIKVFDFERSGL-QHAPEPDAADGPYD 132
SMAD7_HUMAN     RSSYPIFIKSATLDNPDSRT---LLVHKVFPGFSIKAFDYEKAYSLQRPNDHEFMQQPWT 133
                    ***: *  :  *..        * :* *. *: .** .                *:

EmSmadF         GKSLFGPVLHISLGKGWGPAYRRTDFTHCPARLEIWLN     215
SMAD6_HUMAN     -----PNSVRISFAKGWGPCYSRQFITSCPCWLEILLN     165
SMAD7_HUMAN     -----GFTVQISFVKGWGQCYTRQFISSCPCWLEVIFN     166
                        ::**: **** .* *  :: **. **: :*
```

**S6 Figure. Structure and homologies of *Echinococcus* SmadF.** (A) Domain structure of *E. multilocularis* SmadF. Shown is the characteristic DWB (dwarfin B) domain found at the C-terminus of Smads. Note that in contrast to human Smad6 and Smad7, an N-terminal DWA domain is missing in SmadF. Size bar indicates 100 amino acids. (B) Amino acid sequence comparison betwen DWB domains of *E. multilocularis* SmadF, as well as human Smad6 and Smad7. Sittes of perfect alignment (*) as well as groups of strong (:) or weak (.) similarity are marked below the sequences.

S7 Figure

A

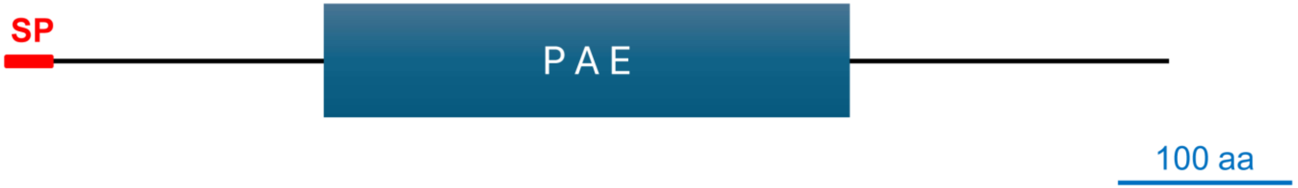


B

```
HsNOTUM              SAQQLNEDLRLHLLLNTSVTCNDGSPAGYYLKESR--GSRRWLLFLEGGWYCFNRENCDS      58
SCHMD_NOTUM          HSRKYQYFQLKKFPNSTNVRCNDGSIPGYYTRPSTTNCSKKWLIFLEGGWYCFNNNTCES      60
EmNOTUM              ------------------------HTDRYYVRRSKDLKSRNWLIYLEGGWYCFDEATCIS      36
                                                 ** : *    *:.**::********:. .* *

HsNOTUM             RYDTMRRLMSSRDWPRTRTGTGILSSQPEENPYWWNANMVFIPYCSSDVWSGASSKSEKN       118
SCHMD_NOTUM         RRRTHYDLFSSEFWSSERQLGGILSNNERINPNFHDYNSVYIPYCSSDLWSGKQLE-KTN       119
EmNOTUM             RQITNHALFSSRTWHMIRYIGGVLSWDDRINPNYHNYNTIFVPYCSSDLWTGKKRE-RTG       95
                    *  *   *:**. *   *   *:** : . ** : : * :::******:*:* . : ...

HsNOTUM             EYAFMGALIIQEVVRELLGR-GLSGAKVLLLAGSSAGGTGVLLNVDRVAEQLEK------       171
SCHMD_NOTUM         GLYFHGSRILDTVVDDLTQNQHFKKVHEVAFVGSSAGGIGVLLNIDRLKRRLKKKLKR--       177
EmNOTUM             GYYFHGSRILTAVIDDLLRQPEFIRVDKVVFSGSSAGGIGVMLNIDRLARRLRRGLRYRR       155
                       * *: *:  *: :*  .  :  .. : : ****** **:**:**: .:*.:

HsNOTUM             -----LGYPAIQVRGLADSGWFLDNKQYRH---TDCVDTITCAPTEAIRRGIRYWNGVVP       223
SCHMD_NOTUM         ---------KVFIHGIVDSAWFLDYPAYRQ---SNCTHIYECPPENALRNGMKLWNPRIP       225
EmNOTUM             WRQSTRNYSRLQISGLVDSAWFLHYPTYAQAYSGSCTSIYDCPPEEGILRGIGLWQPRIP       215
                              : : *:.**.***.   * :    .*.    * * :.: .*:  *:  :*

HsNOTUM             ERCRRQFQEGEEWNCFFGYKVYPTLRCPVFVVQWLFDEAQLTVDNVHLTGQPVQEGLRLY       283
SCHMD_NOTUM         RRCKKFQGRGREWKCFMGPVIYRHLKNPTFIIQSLFDDAQLQMSKVPILEGGSN-KKFSY       284
EmNOTUM             RRCRMAQGKKNFWQCYLGPVIYSHLRTPTFIIQSLFDEAQLQMSRAILLTGGSY-NKLAY       274
                    .**:    . . *:*::*  :*  *: *.*::* ***:*** :... :           *

HsNOTUM             IQNLGRELRHTLKDVPASFAPACLSHEIIIRSHWTDVQVKGTSLPRALHCWDRSLH-         339
SCHMD_NOTUM         IQQLGGFAAQTLRQAKGVFAHSCVDHEILTKSNWAYVSVNNQRLHETLNYWQAYLEG         341
EmNOTUM             IQNLGKAVAKSLDTVQGVFAPACTDHEILTTNHWAQLMVDSQSLHDTLKAWDAHLT-         330
                    **:**    ::*  . . ** :* .***:  .:*: : *..  *  :*: *:  *
```

**S7 Figure. Structure and homologies of *Echinococcus* Notum.** (A) Domain structure of *E. multilocularis* Notum. Shown are the characteristic palmitoleyl-protein carboxylesterase (PAE) domain (PAE) domain and the N-terminal signal peptide. Size bar indicates 100 amino acids. (B) Amino acid sequence comparison betwen PAE domains of Notum from *E. multilocularis* (EmNOTUM), human (HsNOTUM), and Schmidtea mediterranea (SCHMD_NOTUM) origin. Sittes of perfect alignment (*) as well as groups of strong (:) or weak (.) similarity are marked below the sequences.
